# Capture-(Hi)C analysis with PostChicago: Integrative metaprofiling and quantitative comparison of chromatin interactions

**DOI:** 10.64898/2026.10.08.757510

**Authors:** Samuel Krall, Belinda Blum, Valeriia Smialkovska, Ezo Oezcan, Sarah-Victoria Dachtler, Gaia Oldfield, Angelika Feldmann

## Abstract

Functionally linking enhancers to gene promoters at high resolution often entails analysis of 3C datasets, including viewpoint-based methods such as Capture-(Hi)C. Quantitative comparison across samples and experiments, together with visualization and integration of promoter interactions with other datasets are of critical importance. While existing statistical tools identify significant interactions and quantify underlying signal, systematically comparing the shape and extent of this signal across large sets of intervals remains challenging. Moreover, experimental artefacts, such as inefficient crosslinking, can affect downstream interpretation, but are difficult to detect systematically. Here we introduce PostChicago, an R/Bioconductor package that fills this gap. Building upon the widely used interaction-calling package Chicago, PostChicago provides a complete workflow for quality control, visualization, integration and quantitative analysis of promoter interactions between multiple samples, including annotation and quantification of genomic regions based on external datasets. PostChicago extends visual quantitative comparisons beyond individual viewpoints by computing normalized metaprofiles around large sets of intervals that visualize the distribution of interaction signal, enabling comparison between narrow and broad interaction patterns. We further demonstrate how PostChicago can identify experimental crosslinking artefacts. Together, PostChicago enables integrative, shape-sensitive analysis and quality control of promoter Capture-(Hi)C datasets at a scale that would otherwise require extensive custom scripting.

## Introduction

Proximity ligation assays (3C) have emerged as a powerful method to determine interactions between distant sites in chromatin as a proxy of functionally linked gene regulatory regions [1-5]. Capture-(Hi)C is a 3C method for high-resolution and high-sensitivity parallel profiling of chromatin interactions from multiple viewpoints (‘baits’) [6-8]. Statistical analysis of such datasets is enabled by specialized tools such as Chicago [9], CapC-Map [10], ChiCMaxima [11] and CaptureCompendium [12]. Together with software originally developed for 3C/4C analysis (r3Cseq [13] and FourCSeq [14]), these tools provide a statistical framework to determine significant fragment-to-fragment interactions and for initial visualization. peakC [15] and peaky [16] offer alternative detection of significant interactions, whereas other tools, such as Chicdiff [17], have been developed for differential comparison between samples.

A commonality between these tools is that visualization and comparison remain centred on individual viewpoints [10, 12] and/or confined to fragment-level information [9]. What remains missing is the ability to visually and quantitatively integrate the signal across large sets of significant interactions and their genomic surroundings, required to determine the width and structure of interaction domains. This analysis could reveal important principles of genome folding, such as the specificity of the interaction signal at different sites, thereby providing important biological insights. A particular challenge lies in the inherent decay of interaction signal with increasing distance to the bait, making comparisons involving both proximal and distal interactions difficult. Users must therefore currently rely on non-standardized scripts to perform large-scale integration and comparison, which limits reproducibility and comparability between studies.

To address this gap, we developed PostChicago, an R/Bioconductor package operating on the full chicagoData objects produced by Chicago [9], rather than only its significant interaction output. PostChicago provides an integrative reproducible workflow for quality control, visualization, cross-sample comparison and quantification of Capture-C interactions at both restriction fragment and interval (peak) level. In addition to individual viewpoint visualization, significant interactions can be aggregated across large sets of intervals into quantitative metaprofiles, merged by genomic distance into interaction peaks of user-defined width, or quantified under predefined intervals. Because these interactions can be normalized to the peak summit of the reference sample, the extent of signal around an interval set becomes directly comparable between conditions and element classes, allowing narrow and broad interaction domains to be distinguished and quantified. We use new benchmarking datasets and our previously published data to demonstrate two new applications: discrimination of narrow and broad interaction domains during embryonic stem cell (ESC) differentiation and detection of crosslinking artefacts.

## Materials and Methods

### Comparison of interaction domain breadth (Application Example 1)

Metaprofiles were derived and plotted as described in Implementation, using PostChicago v0.99.4. For the formal comparison of interaction widths between interactions overlapping H3K27ac/H3K27me3 and lost/gained interactions in Fig. 2, first the background signal was subtracted from each individual interaction, followed by a subtraction of RA_0h (ESC) signal from the RA_72h signal. For the plot in Fig. 2D and H, the resulting normalised signal of interaction loss was multiplied by -1, for a better comparison with the signal around gained interactions. This normalized differential interaction signal was then plotted as running average signal across k=5 adjacent restriction fragments.

### Fixation perturbation experiment (Application Example 2)

30 Mio culture-grown E14 ESCs (see [18] for tissue culture conditions) were trypsinized, mixed in a falcon tube and split up into three samples (technical replicates) with 10 Mio cells each. Samples were fixed as described previously [18], with Replicate 1 and Replicate 2 fixed with 1.9% formaldehyde (Thermo Fisher Scientific #10751395) and Replicate 3 fixed with 0.95% formaldehyde. Following this, Capture-C libraries were generated using a previously published capture probe pool ([18], Set1) and sequenced with NextSeq 550, as previously described [18].

### Cell culture and retinoic acid differentiation (Application Example 2)

Previously published Capture-C datasets of ESCs prior to and after a time-course of retinoic-acid (RA) treatment were used for all analyses [18]. Only samples captured with the Set1 pool of probes were used.

### Data availability

The high-throughput data reported in this study have been deposited on GEO and are publicly available as of the date of publication. The re-analyzed dataset includes Capture-C (GSE261064, set1) from Mahara et al. [18].

### Software and code availability

PostChicago v0.99.4 was used for all analyses in this study, while data pre-processing was conducted using HiCUP v0.8.3 [19] and CHiCAGO v1.34.0[9], with R version 4.4.1. PostChicago package is available from R/Bioconductor (https://bioconductor.org/packages/devel/bioc/html/PostChicago.html) or GitHub (https://github.com/FeldmannLabDKFZ/PostChicago). Analysis code, including PostChicago source code from v0.99.4 used to generate the figures in this manuscript are deposited on Zenodo (DOI: 10.5281/zenodo.23221805).

### Declaration of generative AI usage in the writing process

Claude 5.5 Opus was used by the authors exclusively to polish and proofread the text in individual passages. No new data, content, or scientific conclusions were generated by this process. The authors reviewed and edited all suggested outputs, assume full authorship accountability, and take absolute responsibility for the accuracy and integrity of the content of this publication.

## Implementation

PostChicago is implemented in R and distributed as an R/Bioconductor package (currently in pre-release). It is integrated with multiple widely used R/Bioconductor packages, including GenomicRanges [20]. The primary input are chicagoData objects containing read counts and scores for all unique pairwise restriction fragment interactions in Capture-C data, which can be generated either using CHiCAGO or via the PostChicago function runChicagoForPostChicago(). Importantly, these objects are similar to bedGraph files, yet differ from the standard CHiCAGO .ibed output, containing only significant interactions. Typical upstream analysis includes HiCUP [19] for read filtering and BAM file conversion to CHiCAGO input (.chinput) files using the CHiCAGO bam2chicago.sh script.

### Data structure and setup

PostChicago analysis is organised around a small number of objects generated from CHiCAGO design files .rmap and .baitmap that are required for the generation of .chinput files. These objects are converted to PostChicago core objects using the helper functions rmap2rmapgr() and baitmap2baited_genes(). A list of chicagoData objects for all samples is generated using the function loadCdList().

### Quality control

Prior to data analysis, plotStats() provides an unbiased quality control step using bait-aligned reads and scores to generate a set of statistical plots, including total numbers of ditags (pairwise fragment-to-fragment interactions), significant interactions and read counts per sample and bait (**Fig. 1 and Fig. 3**). This facilitates the detection of technical artefacts, such as samples with a high total read number but low number of significant interactions, or a shift in read count distributions, such as disproportionately high read counts within significant interactions, indicating the need to overcome high background (see Application 2).

**Figure 1.**
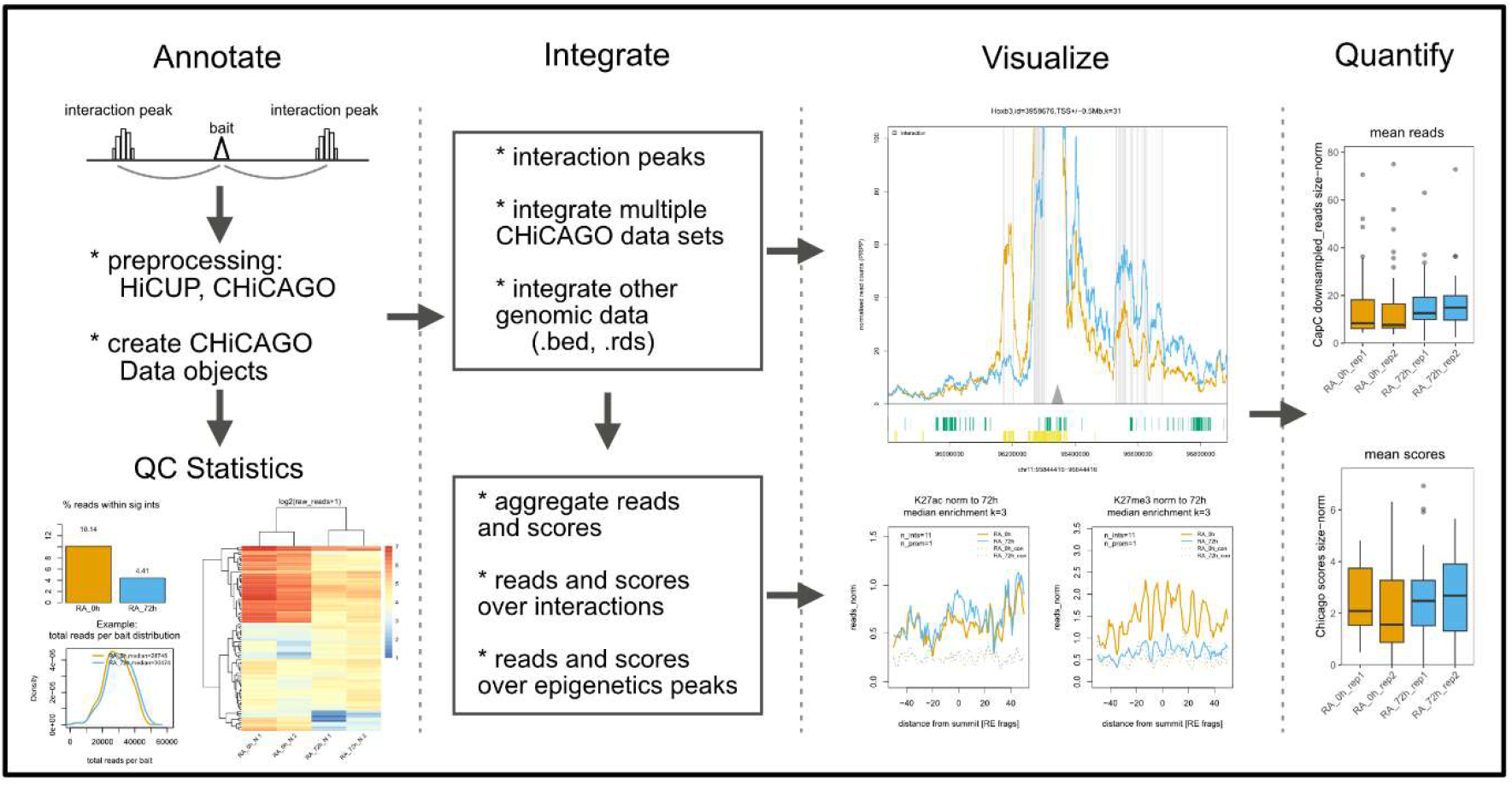
Summary of the PostChicago workflow.

**Figure 2.**
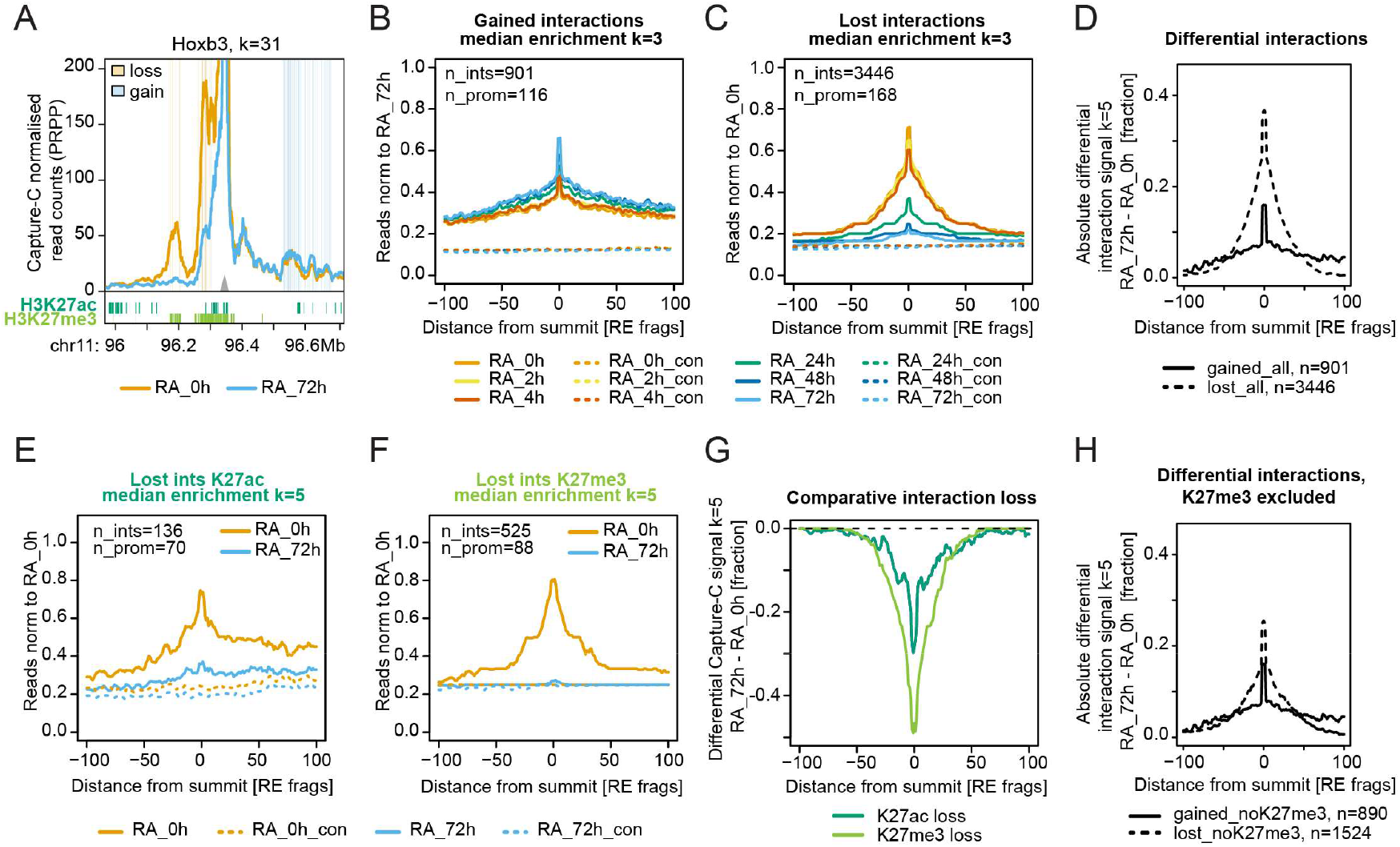
Application 1: Distinguishing narrow and broad interaction domains. **(A)** Example Lineplot generated using plotInteractions() displaying normalized Capture-C signal around the Hoxb3 gene promoter from Mahara et al. [18]. Shown are Hoxb3 interactions before (RA_0h, orange line) and after (RA_72h, blue line) a 72h retinoic acid (RA) treatment of mouse embryonic stem cells. Blue and orange vertical rectangles within the plot represent gained and lost interactions, respectively. H3K27ac (dark green) and H3K27me3 (light green) peak intervals are shown below the plot and correspond to consensus of RA_0h and RA_72h. k: number of DpnII restriction fragments over which the running average signal was calculated. Grey triangle represents the Capture view point (bait). **(B)** Normalized metaprofiles produced using getMatrix() and plotAggregatePeaks() and displaying the interaction signal centred at gained interactions across the differentiation time course (0, 2, 4, 24, 48, 72h RA treatment). Average signal was calculated across 901 gained interactions by a total of 116 promoters (baits). Dotted lines: Control Capture-C distance-matched signal at the same baits. Reads were normalized to the central fragment in the RA_72h sample and plotted as the running average across k=3 restriction fragments. Note the focal gain of signal around the central fragment and elevated signal extending to +/-100 restriction fragments. **(C)** As in B, but for lost interactions (total: 3446, captured by 168 view points) and upon normalization to the RA_0h sample. Different reference sample from (B) was chosen in order to ensure normalization to a positive control, providing a less noisy signal. Note the overall broader central signal distribution and near complete reduction of the signal upon differentiation and towards +/-100 restriction fragments. **(D)** Normalized absolute differential Capture-C signal between RA_72h and RA_0h for gained (black line) and lost (dotted line) interactions. Signal was first normalized to the background signal via subtraction, then RA_0h were subtracted from RA_72h. For a better comparison of the signal width, lost interaction signal was multiplied by -1. Plotted is the average signal across k=5 restriction fragments. Note the wider peak around lost (dotted line) than gained (solid line) interactions and an elevation of gained interaction reads towards +100 restriction fragments. The asymmetry likely reflects the directionality of the metaprofiles (+100 is closer to the bait) and an overall stronger promoter proximity of interaction gain. **(E)** As in B, but metaprofiles are centred at H3K27ac peaks overlapping lost interactions. Shown is only the signal before (RA_0h) and after (RA_72h) a 72-hour RA treatment. The plots were generated using makeOneGeneOnePeak(), followed by getMatrix() and plotAggregatePeaks(). k: number of DpnII restriction fragments over which the running average signal was calculated. Average signal was calculated across 136 lost interactions by a total of 70 promoters (baits). Dotted lines: Control distance-matched Capture-C signal at the same baits. Reads were normalized to the central fragment in the RA_0h sample and plotted as running average across k=5 restriction fragments. Note the increase towards +100 restriction fragment representing the increase of signal towards the bait. **(F)** As in (E) but for H3K27me3 peaks, over 525 interactions and 88 promoters. **(G)** Normalized differential Capture-C signal between RA_72h and RA_0h for H3K27ac (dark green) and H3K27me3 (light green) peaks overlapping lost interactions. Signal was first normalized to the background signal via subtraction, then RA_0h were subtracted from RA_72h. Plotted is the average signal across k=5 restriction fragments. Note the wider peak around H3K27me3 than H3K27ac-centred interactions. **(H)** As in (D), but excluding all H3K27me3-overlapping interactions for both gained and lost (dotted line) interactions.

**Figure 3.**
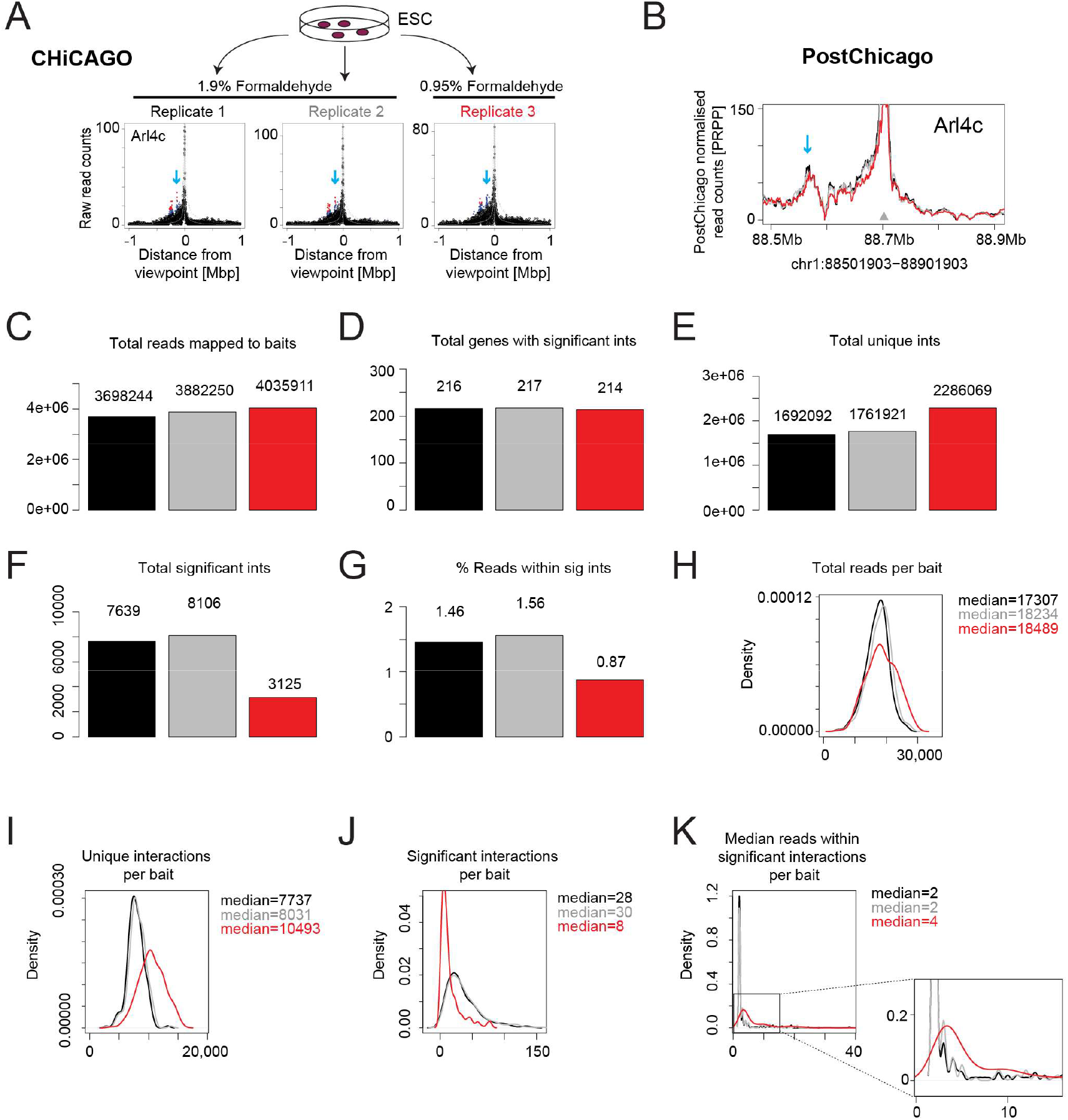
Application 2: Detection of Crosslinking artefacts by PostChicago. **(A)** Schematic: Three chromatin samples derived from a single mouse embryonic stem cell (ESC) culture were crosslinked with either 1.9% (Replicate 1 and Replicate 2, black and grey, respectively) or 0.95% (Replicate 3, red) formaldehyde. Chicago output scatter plots show enrichment +/-1Mb around the Arl4c gene promoter. Red: significant fragment-to-fragment interactions. Blue arrow: highlights an interaction ∼150kb from the bait. (**B**) Lineplots generated using plotInteractions() displaying normalized running average Capture-C profiles of Arl4c. Blue arrow: highlights an interaction ∼150kb from the bait matching the interaction highlight by the blue arrow in (A). (**C-K**) Output of plotStats(), including (**C**) total read counts captured by baits, (**D**) the total number of genes with significant interactions, total unique (**E**) and significant (**F**) interactions. (**H-K**) Display distributions of total reads (**H**), unique (**I**) and significant (**J**) interactions per bait. (**K**) Per bait distribution of median read number within significant interactions. Black: Replicate 1 (1.9% Formaldehyde), grey: Replicate 2 (1.9% Formaldehyde), red: Replicate 3 (0.95% formaldehyde).

### Visualization

PostChicago’s core visualization function, plotInteractions(), integrates data from the list of chicagoData objects to produce normalized line plots, which display the running mean of normalized read counts over k adjacent fragments and across a user-defined genomic window surrounding the bait (zoom) or over exact intervals (xlim). Interaction signal is further normalized to the sequencing depth of bait-interacting fragments per 100,000 reads and to the total number of baits within a single Capture-C experiment, enabling a comparison between experiments with different capture designs. The resulting plots can be annotated with genomic intervals (for instance ChIP-Seq peaks loaded from .rds or .bed files using loadGrl()), or significant interactions (see Data integration) as shaded rectangles with optional colouring for different types of interactions (e.g. lost and gained). Additional flexibility is ensured by the generation of sample-normalized BedGraph files for each bait by makeBedGraphs(), which also outputs BED files with bait location and the locations of significant interactions to facilitate exploration in genome browser.

### Data integration

makeIntsTable() integrates multiple data sets by extracting all significant fragment-to-fragment interactions across a set of samples from one Capture-C experiment, while assigning a unique interaction identifier intID. These significant interactions are populated with quantitative information, such as raw and downsampled weighted average and replicate read counts, as well as CHiCAGO scores per sample. For scalability, processing is chunked depending on the total number of baits in an experiment. Interactions can be annotated with genomic intervals (e.g. ChIP-Seq peaks) by annotateInts(), where the overlap distance is controllable in a user-defined manner by the maxgp parameter. makeQCplots() then provides sample-to-sample correlations and hierarchical clustering of statistically significant interactions, necessary for the assessment of reproducibility and batch effects.

### Quantification within and around intervals of interest

Epigenetic peaks can span a range of 300-10,000bp, often covering multiple restriction fragments, with a possibility of multiple significant interactions with a single viewpoint. For quantitative integration with intervals, PostChicago therefore first generates tables connecting each interval only once with each viewpoint using the function makeOneGeneOnePeak(). quantifyWithinPeaks() annotates oneGeneOnePeak tables with read and score coverage within the entire interval, including the signal within not statistically significant fragment-to-fragment interactions. boxplotsCapC() function provides boxplot summaries of this interval-level quantification.

Similar to ChIP-seq peaks, interaction peaks can contain several statistically significant interactions. Two additional PostChicago functions offer the possibility to summarize individual fragment-to-fragment interactions to interaction peaks based on fusing multiple interactions by either inter-fragment distances (aggregatePeaks()) or distances in base pairs (aggregatePeaks_regions()). The total read/score coverage of interaction ‘peaks’ is then quantified by quantifyWithinPeaks() and visualized by boxplotsCapC() analogous to oneGeneOnePeak tables.

Visualization of the interaction signal surrounding statistically significant interactions, interaction peaks or intervals is facilitated by getMatrix(), which identifies the central fragment within the distal site interacting with viewpoints and computes interaction matrices centring on these fragments and extending for a predefined distance in restriction fragments. Since Capture-C signal naturally increases towards viewpoint proximity, the matrices are automatically stranded in a way that the viewpoint is always to the right of the distal site. Finally, for each interaction, a distance-matched control is extracted, based on the interaction signal on the other side of the viewpoint.

In addition to plotting downsampled read counts, getMatrix() can be carried out in the ‘scaling’ mode, where reads are plotted based on a normalization to the total number of viewpoints and library depth per 100,000 mapped reads. This normalization recapitulates the normalization provided by the plotInteractions() function. Depending on the sequencing experiment, this can result in a stronger signal, but should generally not affect inter-sample comparison. Another optional normalisation provides a means to ensure that the observed signal is not driven by promoter-proximal interactions, which inherently display higher read counts than more distal interactions. To normalize for these differences, each row in the matrix (displaying interaction signal around the central interaction fragment) is normalized to the interaction signal at the central fragment of a reference sample (typically positive control). For each viewpoint/distal site pair, this expresses the surrounding signal relative to the summit of the reference sample, making profiles directly comparable between samples.

The output of getMatrix() can next be plotted as metaprofiles via plotAggregatePeaks(), enabling a visual comparison between different samples at and surrounding focal interaction points.

A fully worked example of the complete workflow, including the functions not featured in the applications below, is provided in the package vignette.

## Results

### Application 1: Distinguishing narrow and broad interaction domains

Cell-type transitions are accompanied by alterations in genome topology, including extensive loss and gain of promoter interactions [18, 21-27]. Although the relationship between differential interactions, linked chromatin modifications, and transcription have been extensively studied, it remains unclear whether there are fundamental distinctions in the shape and structure of dynamic interactions.

We used PostChicago to compare the topological structure of dynamically gained and lost interactions during retinoic-acid(RA)-dependent ESC differentiation in our previously published datasets [18]. Initial visual inspection of comparative line plots around the Hoxb3 gene promoter produced by plotInteractions() suggested differences in the breadth of lost and gained interactions and the chromatin structure of the surrounding chromatin (**Fig. 2A**). To quantitatively compare these differences across all lost (n=3446) and all gained (n=901) interactions, we computed interaction matrices centred at either lost or gained interactions and extending 100 restriction fragments to either site using getMatrix(). We next plotted normalized interaction metaprofiles by plotAggregatePeaks(), revealing that interaction gain occurs focally, within a region with generally increased interaction density (**Fig. 2B**). In contrast, loss of interactions extended over a broader domain, reaching near background levels upon 72 hours of RA treatment (**Fig. 2C**). This broader structure of interaction loss (**Fig. 2D**) was observed at a range of distances, confirming that the difference in shape is not explained by differences in the distance distributions between lost and gained interactions (**Fig. S1A-B**).

Loss of interactions during RA-mediated ESC differentiation preferentially occurs at sites overlapping H3K27me3 peaks, whereas gain of interactions displays a higher overlap with H3K27ac [18] (**Fig. 2A**). Since H3K27me3 peaks are part of Polycomb domains, spanning large regions in ESCs [28], we considered the possibility that the increased breadth of lost interactions is driven by their association with Polycomb sites. To test this, we first computed interaction matrices centred at either H3K27me3 or H3K27ac peaks, overlapped them with distal sites whose interactions were lost during the differentiation, and plotted the corresponding metaprofiles. As expected, more H3K27me3 peaks (n=525) overlapped lost interactions than H3K27ac peaks (n=136), with interaction loss around H3K27me3 peaks spanning visibly broader regions than those overlapping H3K27ac peaks (**Fig. 2E-G**). This confirms that H3K27me3-overlapping interactions are associated with a broader loss than H3K27ac-overlapping interactions. To compare the general properties of interaction gain and loss, not confounded by the association with H3K27me3, we removed all H3K27me3-overlapping interactions from the entire set of lost interactions and replotted the corresponding metaprofiles. Interestingly, even when uncoupled from H3K27me3, interaction loss still occurred over broader domains than interaction gain (**Fig. 2H** and **Fig. S2D**), although now with a lower magnitude than when H3K27me3-overlapping interactions were included (26% loss vs. 38% loss) (**Fig. 2D** and **Fig. S1C**). Thus, loss of interactions is characterized by particularly broad domains, with H3K27me3-associated loss being particularly strong. On the other hand, gain of interactions appears within regions of already elevated overall interaction densities.

### Application 2: Detection of crosslinking artefacts by quality control metrics

Inadvertent experimental artefacts can arise during Capture-(Hi)C library preparation, including reduced crosslinking, digestion or ligation efficiencies. While each of these can directly affect the detection of significant interactions and biological interpretation of the data, chromatin crosslinking artefacts are particularly difficult to distinguish from normal variation between high-quality samples. We reasoned that under-crosslinking should theoretically result in higher background, as well as in lower numbers of significant interactions. Since part of the quality control pipeline provided by PostChicago’s function plotStats() is comparing quantitative metrics across samples, we hypothesized that these statistics should be useful in detecting under-crosslinked samples in a quantitative manner.

To experimentally test the ability of PostChicago to detect crosslinking artefacts, we crosslinked samples derived from the same ESC culture with either 1.9% (standard) or 0.95% (half) concentrated formaldehyde (**Fig. 3A**), keeping the remaining steps of library preparation consistent across samples. Visualization of interactions at an individual view point (Arl4c) using Chicago indeed revealed elevated background and a reduced number of significant interactions (**Fig. 3A**). Yet, given the observable variation between the samples crosslinked with standard amounts of formaldehyde (Replicates 1 and 2), the background increase was not clearly discernible as significantly elevated, and significant interactions were still detectable at expected positions. In contrast to Chicago, the plotInteractions() function of PostChicago, which uses inter-sample normalization of read counts, flattened the background (**Fig. 3B**), however was still able to detect a small decrease in read count enrichments within a significant interaction localized approximately 130kb upstream of the Arl4c gene promoter. We conclude that at this locus, visual inspection did not clearly separate under-crosslinking from normal variation between replicates.

We next tested whether PostChicago’s quality control diagnostics provided by plotStats() are able to close this gap. Upon sequencing, all three samples displayed a similar number of informative reads and captured genes (**Fig. 3C-D**). As expected for reduced crosslinking and correspondingly increased background, unique interaction number increased in the under-crosslinked sample (**Fig. 3E** and **Fig. 3H-I**), while the total number of significant interactions decreased by at least 60% (**Fig. 3F**). This was not driven by distinct overall numbers of sequenced reads, as the total number of reads mapping to baits were comparable between samples. While significant interactions per bait were reduced from 28-30 to 8 (**Fig. 3J**), the most striking indication of crosslinking artefacts was visible when comparing the median number of read counts within a significant interaction (**Fig. 3K**). Here, reduced crosslinking resulted in a strong shift of the distribution towards a higher number per interaction, consistent with a requirement of higher read counts to overcome elevated sample background.

Importantly, shifts in read counts can also be observed under biological conditions, when comparing Capture-C samples with an expected increase in read counts. For example, this was evident when we compared read counts within a published Capture-C dataset that was biased for genes induced during ESC differentiation [18] (**Fig. S2**). These samples displayed shifts corresponding to under-crosslinking (**Fig. S2A-B**), including increased numbers of reads per significant interaction upon differentiation (**Fig. S2C**). Of note, there is a clear distinction between this biological difference and crosslinking artefacts. Higher read count within significant interactions of induced genes (**Fig. S2**) presents as a longer interaction tail (or bimodal distribution when plotted on a log scale), suggesting that this increased read count is not an artefactual feature of interactions in differentiated cells, but rather a true increase in interaction intensities that accompanies gene activation at a subset of sites. This differs from the global shift in read counts observed at significant interactions in under-crosslinked samples (**Fig. 3K**). Reduction in total read counts in these samples can be explained by an extensive global interaction loss observed upon ESC differentiation [18].

Taken together, reduced crosslinking produced a characteristic shift during the calculation of quality control metrics. Because a similar shift can arise biologically at induced genes (**Fig. S2**), the signature is interpretable as a technical artefact when it is restricted to a subset of replicates and/or displays a global, unexpected shift.

These examples demonstrate how PostChicago can be used to identify broad and narrow interaction domains and provide biological insights into the reprogramming of topologies during ESC differentiation.

## Discussion

In this study we present PostChicago, an R/Bioconductor package designed as a versatile toolkit for Capture-(Hi)C analysis downstream of available callers of significant interactions. Our package facilitates biological inference by providing a full workflow of functions for integration, annotation, visualization and comparative quantitative analysis of Capture-C datasets, thereby complementing and combining the capabilities of similar tools [9, 10, 12, 17] in a single R package. We demonstrate PostChicago’s two distinguishing capabilities: The ability to quantitatively compare the width and shape of distinct types of interactions and their surroundings through meta-profiling, and its ability to detect experimental crosslinking artefacts.

In a key application, we use PostChicago to show that during RA-mediated differentiation of ESCs, interaction gain is more focal than interaction loss. Interestingly, gain of interactions occurs within regions of interaction density already elevated in ESCs (**Fig. 2**), consistent with the possibility that elevated interactions (such as those established within TADs by cohesin [29-31]) may prime chromatin for the creation of specific enhancer-promoter interactions [32, 33]. In contrast, interaction loss occurs over broader domains, but results in complete loss of promoter interactions. This effect is independent of the distance over which interactions occur, suggesting that a broader profile may be a general property of interaction loss. On the other hand, loss of Polycomb-associated interactions generally occurs over broader domains than loss of H3K27ac-associated interactions, compatible with the idea that chromatin modifications shape dynamic interaction profiles.

Moreover, we show that PostChicago can also detect crosslinking artefacts, extending existing quality control tools [12] to the specific detection of crosslinking artefacts, which has previously required inspection of individual loci. We demonstrate this ability using ratios between the number of significant interactions called and the total read number within significant interactions (**Fig. 3**).

Taken together, apart from providing quality control possibilities that, to our knowledge, are not available in existing Capture-C analysis tools, we show that PostChicago’s ability to aggregate large numbers of interactions to metaprofiles can facilitate genuine biological discoveries related to size and shape of interaction domains.

### Limitations

A limitation of the workflow is that currently PostChicago relies on the output of Chicago, a very specific interaction caller package. Future versions will adapt its usage to other available peak calling approaches [10, 11, 13-16]. We have shown the capability of PostChicago to identify under-crosslinking artefacts of known high degree of under-crosslinking (50%). It remains to be shown whether detecting artefacts with a lower under-crosslinking is more challenging.

## Author Contributions

A.F. designed the study. A.F., S.K., B.B. and S.V.D. wrote the code for the package. S.K. and A.F. created the R/Bioconductor package. S.K. and A.F. wrote the original manuscript. S.K. B.B., V.S., G.O., E.O., S.V.D., A.F. wrote the manuscript. B.B. and A.F. made the figures. S.K. and A.F. analysed the data. B.B., V.S., G.O., E.O. and S.V.D. tested the package.

## Acknowledgements

We thank the members of the Feldmann lab for their input during the testing phase of the package and for the critical proofreading of the manuscript.

## Funding

Work in the Feldmann lab is supported by the European Research Council (Award Number: ERC-STG 101115812), the Helmholtz Association (Helmholtz Young Investigator Award Number: VH-NG-1604), and the Health + Life Science Alliance Heidelberg Mannheim.

---

**Figure S1.**
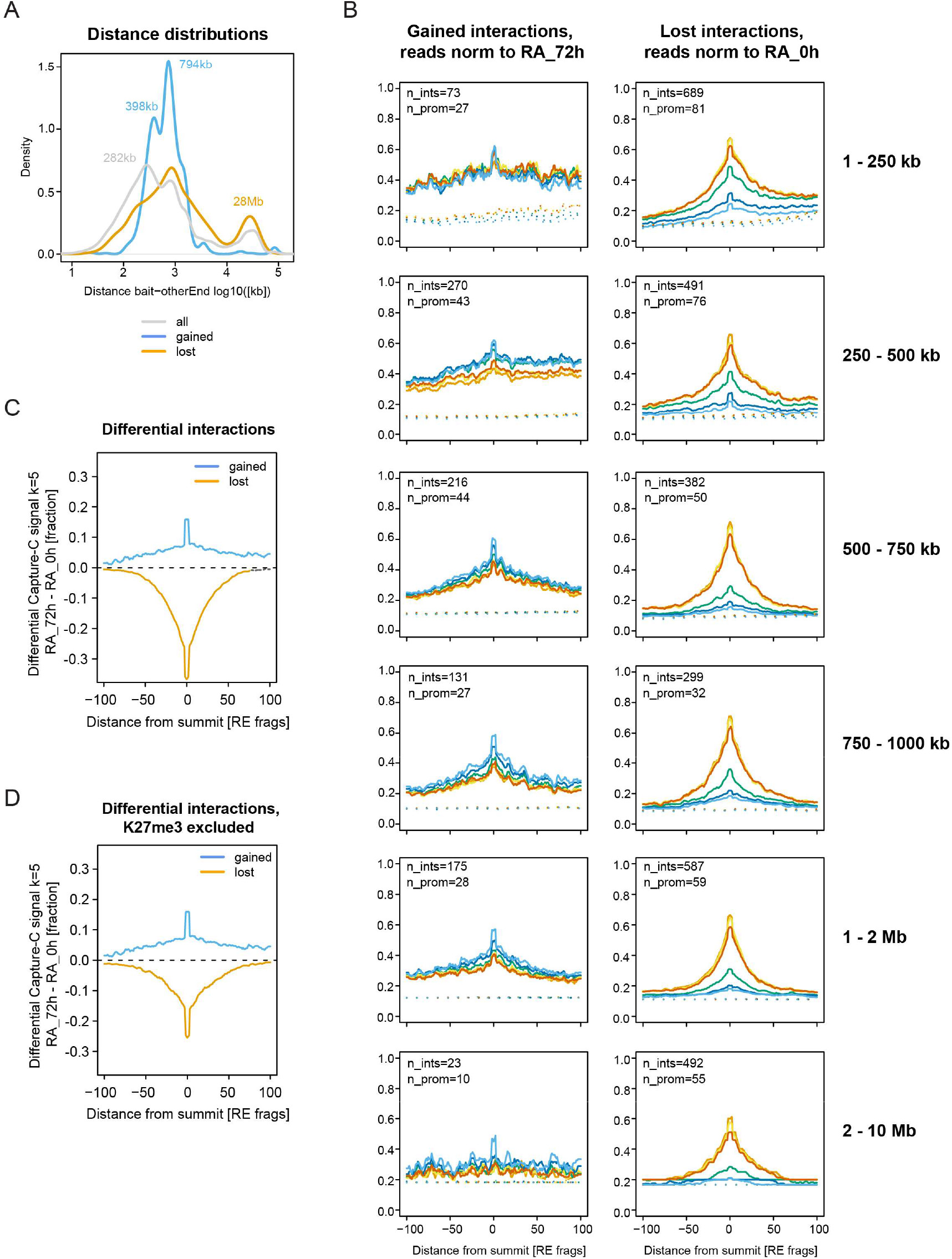
Application1: Distinguishing narrow and broad interaction domains. **(A)** Distance distributions (log10) of lost (orange), gained (blue) and all interactions. Note the bi-to tri-modal distribution of distances. The approximate distances are displayed at the distribution summits, with gained interactions enriched around 400kb and 800kb, whereas lost interactions show summits around 800kb and 28Mb. **(B)** Normalized metaprofiles produced using getMatrix() and plotAggregatePeaks() displaying the interaction signal centred at gained (left) or lost (right) interactions across the differentiation time course (0, 2, 4, 24, 48, 74h RA treatment). Signal was stratified by distances, as displayed on the right. The total number of interactions and promoters across which the average signal was calculated are displayed within each plot. Dotted lines: Control Capture-C distance-matched signal across the same baits. Reads were normalized to the central fragment in the RA_72h sample and plotted as running average across k=5 restriction fragments. **(C)** Normalized absolute differential Capture-C signal between RA_72h and RA_0h for gained (black line) and lost (dotted line) interactions. Signal was first normalized to the background signal via subtraction, then RA_0h were subtracted from RA_72h. Plotted is the average signal across k=5 restriction fragments. Note the wider peak around lost (dotted line) than gained interactions and an elevation of gained interaction reads towards 100 restriction fragments. The asymmetry likely reflects the directionality of the metaprofiles (+100 is closer to the bait) and an overall stronger promoter proximity of interaction gain. **(D)** As in (C), but excluding all K27me3-overlapping interactions for both gained and lost (dotted line) interactions.

**Figure S2.**
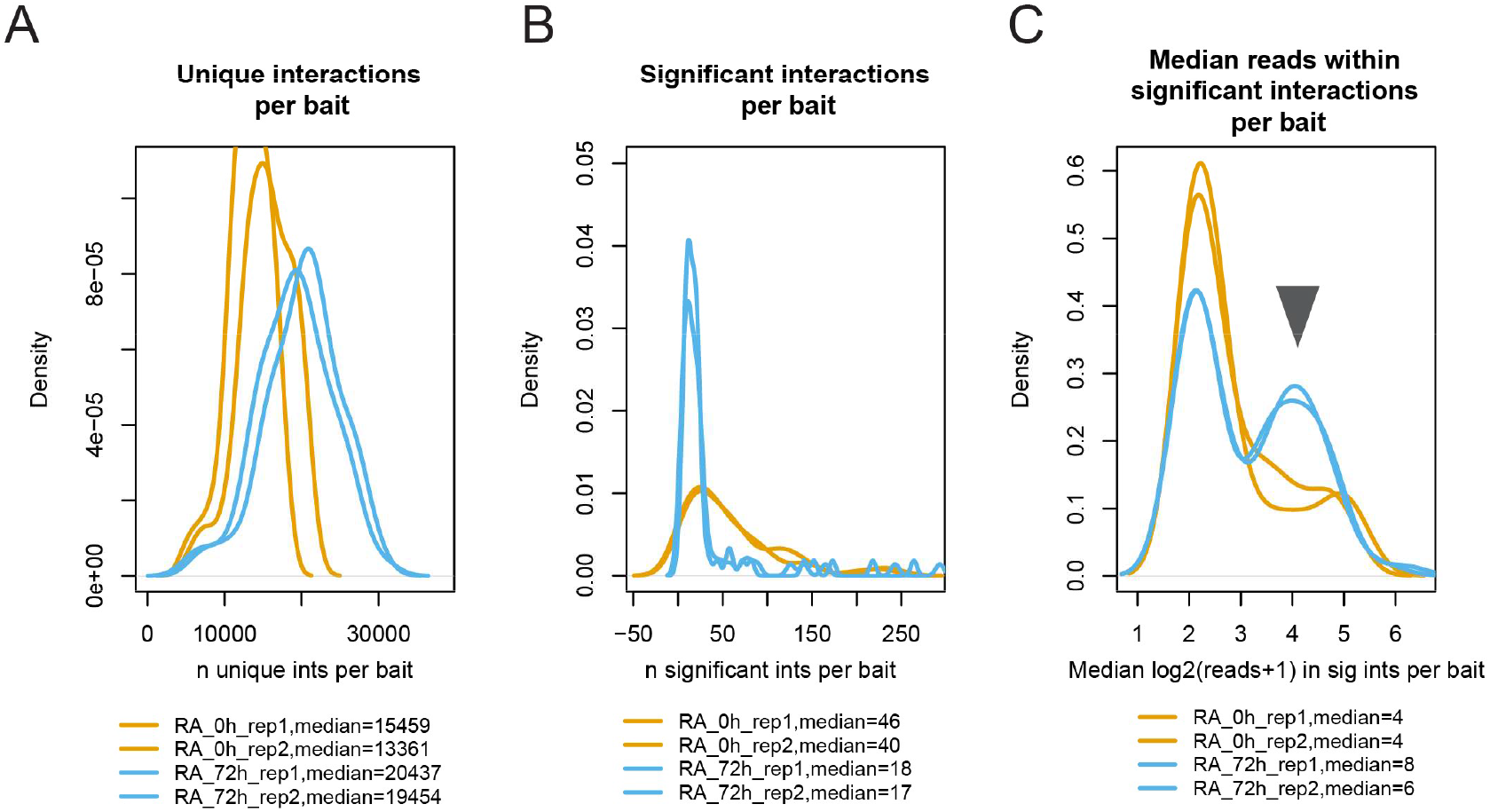
Application2: Detection of Crosslinking artefacts by PostChicago for induced genes. Shown are plots derived using plotStats() from published Capture-C data [18], focusing only on induced genes. Shown are the distributions of unique (**A**) and significant (**B**) interactions per bait, as well as the per bait distribution of median read number within significant interactions (**C**). Grey triangle in C points to the shift in distribution visible as one of two distribution peaks in RA_72h samples. Orange: Samples prior to RA treatment. Blue: Samples after a 72h RA treatment.

## Notes

### Competing Interest Statement

The authors have declared no competing interest.

